# Osmoregulation of glutamine synthetase from Giant freshwater prawn (*Macrobrachium rosenbergii*) under osmotic stress

**DOI:** 10.1101/517409

**Authors:** Zhijie Lu, Zhendong Qin, V Sarath Babu, Chengkai Ye, Guomao Su, Jiabo Li, Guang Yang, Haiyang Shen, Gan Pan, Li Lin

**Author notes:** Corresponding author: College of Animal Sciences and Technology, Zhongkai University of Agriculture and Engineering, Guangzhou, Guangdong, 510225, China, Li Lin:; Guangdong Provincial Key Laboratory for Healthy and Safe Aquaculture, College of Life Science, South China Normal University, Guangzhou 510631, China, Gan Pan.

## Abstract

Glutamine synthetase is a key enzyme that catalyzes the biosynthesis of glutamine (Gln) from glutamate and ammonia. Gln a vital amino acid acts as a precursor for protein synthesis and also assist in ammonia repressor and a key osmoregulators in aquatics. Here, we report the cloning and characterization of the GS gene from *Macrobrachium rosenbergii* (*Mr*-GS). The complete nucleotide and deduced amino acid sequences were determined that phylogenetically shared highest identity with other crustaceans. GS mRNA was differentially expressed in 6 different tissues, with high to low order as muscle > gills > heart > stomach > brain > haemolymph. *Mr*-GS expression and the glutamine concentrations were analyzed in the gills and muscle tissues of prawn under hyper/hypo-osmotic stress conditions. Under hyper-osmotic stress, the mRNA expression of *Mr*-GS was significantly increased in both gills and muscle at 3, 6 and 12 h post-treatment with 2.54, 4.21 and 10.83 folds, and 11.66, 17.97 and 45.92 folds, respectively. Protein analysis by western blot (WB) and Immunohistochemistry (IHC) further confirmed the *Mr*-GS expression was increased at 12 h post treatment. On the other hand, under hypo-osmotic stress, the mRNA expression of *Mr*-GS was also significantly increased in both gills and muscle at 3, 6 and 12 h post treatment with 1.63, 3.30 and 3.52 folds, and 4.06, 42.99 and 26.69 folds, respectively. Furthermore, under hyperosmotic stress, Gln concentration was increased in both gills and muscle at 6 and 12 h post treatment with 1.83, 2.02 folds, and 1.41, 1.29 folds, respectively. While, under hypo-osmotic stress, Gln concentration was increased in both gills and muscle at 3, 6 and 12 h post treatment with 3.99, 3.40, 2.59 folds, and 1.72, 1.83, 1.80 folds, respectively. Taken together, these results suggest that *Mr-*GS might play a key role in osmoregulation in *M*. *rosenbergii*.

## Introduction

Giant freshwater prawn (*Macrobrachium rosenbergii*) is one of the world’s largest freshwater cultured crustaceans and has a wide distribution in tropical and subtropical areas of the world [1, 2]. There have been reported that the prawn could mature and spawn in the freshwater area [3]. However, they must migrate to the brackish water with salinity range between 9-19 ‰ for hatching and nursing of the larvae [2, 4]. As a result of migration, this species exhibits an excellent tolerance to a wide range of salinity, which is a characteristic of the prawn [5]. There are a number of reports about the salinity tolerance of the prawn [2, 4-10]. However, the mechanism underlying the osmoregulation of the prawn remains enigmatic.

Glutamine synthetase (GS, EC 6.3.1.2) is an enzyme catalyzes a reaction that incorporates ammonium into glutamate and generates Glutamine (Gln), i.e., Glutamate + ATP + NH_3_ → Glutamine + ADP + phosphate [11]. The Gln plays crucial roles in an array of biochemical functions, including protein synthesis, lipid synthesis, cell growth, energy supply, as well as ammonia carrier [12]. The GS gene has been reported in many species which included not only vertebrate species such as Chinese hamster [13], chicken [14] and human [15], but also invertebrate species like *Procambarus clarkii* [16], *Crassostrea gigas* [17], *Fenneropenaeus chinensis* [12] and *Litopenaeus vannamei* [18]. However, up to date, there is no information about the GS gene of *M*. *rosenbergii*. Previous studies had mainly focused on salinity-related changes in oxygen consumption, ammonia excretion, and ion osmoregulation in *M*. *rosenbergii* [5]. There have been reported that with the increase of water salinity, the levels of some free amino acid (FAA), including glycine, proline, arginine, glutamate, and alanine in the tissues and haemolymph also raised in *M*. *rosenbergii* and *M*. *nipponense* species [19, 20], due to the catabolism of proteins and amino acids (AA) [21]. Ammonia which is toxic for aquatic animals and it must be catalyzed into Gln with the effort of GS enzyme as a nontoxic transporter in the haemolymph circulation [22].

Once the crustaceans were stressed by various environmental factors, GS expression and the concentration of Gln were found to be increased, illustrating that GS plays an essential role in environmental stress resistance and adaptation [12, 23]. Furthermore, Gln as a transporter of ammonia in the haemolymph provides an abundant FAA as osmolytes that further utilized by other cells or protein synthesis [18, 20]. Since the gills is a vital tissue for osmoregulation and the muscle is the largest storehouse of protein and amino acid for providing energy, therefore, we focused our studies on gills and muscle in this report. We have cloned a GS gene from *M*. *rosenbergii* (*Mr*-GS), and the osmoregulation of the GS was characterized under osmotic stress.

## Materials and methods

### Experimental design and samples collection

Adult *M*. *rosenbergii* (approximately 12-15 g) were obtained from Jin Yang Aquaculture Co. Ltd., Guangzhou, China. First of all, the prawns were acclimatized to freshwater in the tank at 25 °C for at least one week before experiments. The prawns were cultured in freshwater and used as negative control. For hyper-osmotic stress treatment, some of the prawns were shifted directly into brackish water with 13 ‰ salinity. For hypo-osmotic stress treatment, the prawns which have been adapted to the water with 13 ‰ were shifted directly to freshwater. The tissues from six individuals were sampled for the RNA extraction at 0, 3, 6, and 12 h post the stress treatment. The samples of the gills and muscle were frozen immediately in liquid nitrogen for western-blot analysis, while part of them was instantly fixed and processed for immunohistochemistry (IHC).

### RNA isolation, cDNA synthesis and gene cloning of Mr-GS

The total RNA from the various samples were extracted using RNAiso plus (TaKaRa, Dalian, China) and the first-strand cDNA was synthesized using HIScript^®^ Q Select RT SuperMix for qPCR (Vazyme, Nanjing, China) according to the manufacturer’s instructions. Specific primers was designed based on the sequences of the GS gene identified from the *M*. *rosenbergii* transcriptomic data in our laboratory (unpublished data), so as to amplify the complete open reading frame ORF of *Mr*-GS. All used primers were shown in Table 1. PCR amplification was performed under the following conditions: pre-denaturation at 95 °C for 5 min, followed by 35 cycles of 95 °C for 10 s, 55 °C for 30 s and 72 °C for 1 min, post extension at 72 °C for 10-min and finally kept at 4 °C. The amplified specific PCR products were electrophoresed on 1% agarose gels, and the target products were purified with a TaKaRa Agarose Gel DNA Purification KitVer.2.0 (TaKaRa, Japan). The purified DNA fragments were ligated into the pET-32a (+) plasmid (TaKaRa, Japan) and transformed into competent *Escherichia coli* DH5α cells. Positive clones containing inserts of the expected size were sequenced using M13 primers and sequenced at Invitrogen, Shanghai.

**Table 1.**
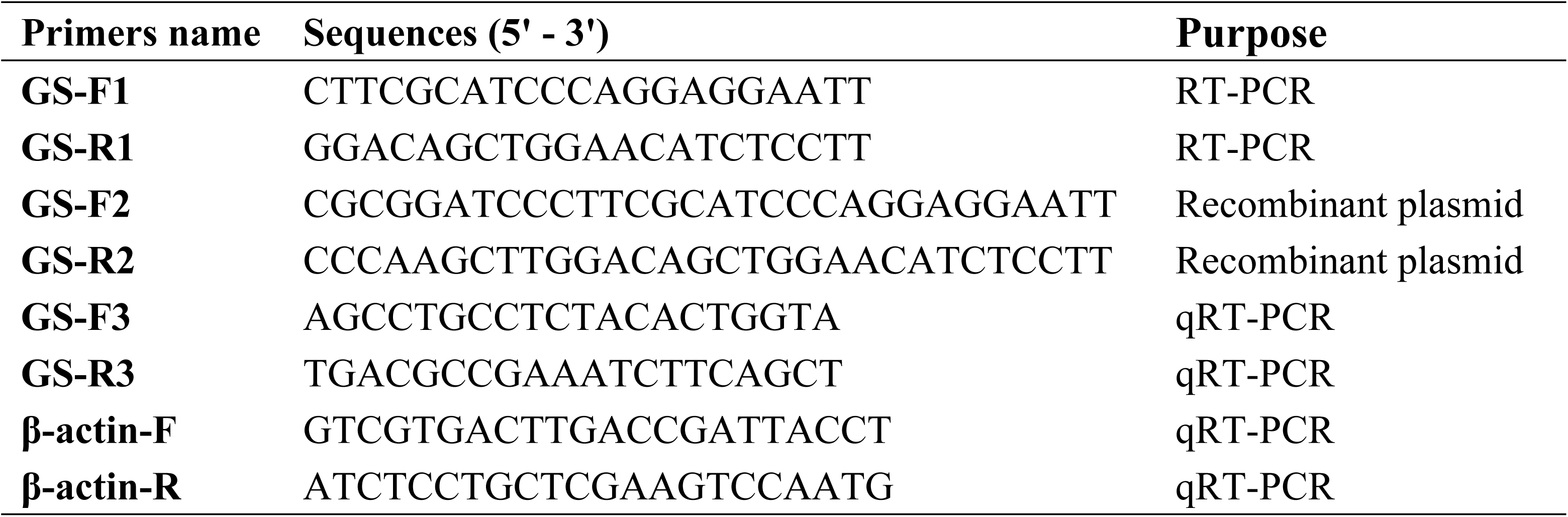
Primers used in the present study

### Sequence and phylogenetic analysis of Mr-GS

The full-length AA sequence of *Mr*-GS ORF and protein domains were predicted by Emboss (http://emboss.Bioinformatics/) and SMART (http://smart.embl-heidelberg.de) tools. The sequences similarity was analyzed by the BLAST program (http://www.ncbi.nlm.nih.gov/blast). Multiple sequence alignments were performed by Clustal X 2.0 program, and the breakpoints analyses were further determined by DNAMAN software package (Lynnon Biosoft, Canada). The phylogenetic tree was constructed based on the ORF AA sequences of *Mr*-GS proteins by MEGA 6.0 software with the neighbor-joining (NJ) method with 1000 bootstraps replications.

### Quantitative real-time PCR (qRT-PCR) assay and data analysis

The expression pattern of *Mr*-GS in various tissues at different time points was studied using qRT-PCR in Roche LightCycler 480 (Roche, USA). All used primers were presented in Table 1, where β-actin was used as an internal reference gene. The qRT-PCR was conducted using AceQ^®^ qPCR SYBR^®^ Green Master Mix (Vazyme, Nanjing, China). The reaction was performed in a final volume of 20 μl, containing 1 μl cDNA, 10 μl AceQ^®^ qPCR SYBR^®^ Green Master Mix, 1 μl each specific primer, and 7 μl ddH_2_O under following conditions: 95 °C for 3 min, followed by 40 cycles of 95 °C for 15 s, 60 °C for 30 s and 72 °C for 20 s, finally at 4 °C for 5 min on. The relative expression ratio of the target genes versus β-actin gene was calculated using 2^-ΔΔCT^ method [24]. Each sample was measured at least triplicate, and all data were presented as mean ± standard deviation (SD). Significant differences between samples were analyzed by one-way analysis of variance (ANOVA) in GraphPad Prism 7. The difference was considered significant, *P* < 0.05 (*), *P* < 0.01 (**) or not significant, *P* > 0.05 (NS).

### Western-blot assay

Total protein from the lysates of frozen gills and muscle tissues were prepared by homogenization as described [25]. Briefly, the tissue samples were weighed and homogenized three times in 5 volumes (w/v) of ice-cold extraction buffer containing 50 mM imidazole (pH 7.0), 1 mM EDTA, 25 mM NaF, and 1 mM PMSF. Subsequently, sonicated for 30 s and centrifuged at 10,000 × *g* at 4 °C for 10 min. The protein concentrations were determined according to the method of Bradford Protein Assay Kit (Beyotime, Shanghai, China). The total protein (about 50 μg) was separated in an SDS-PAGE (10 %) and transferred onto nitrocellulose membrane (Bio-Rad, America). The membranes were blocked with TBST (137 mM NaCl, 20 mM Tris, 1 % Tween-20, pH 7.6) containing 5 % skim fat milk at room temperature (RT) for 1 h. Then the membranes were incubated at 4 °C overnight with the rabbit *anti*-*Mr*-GS (1/1000 dilution) primary antibody which was prepared in our laboratory. Subsequently, the membranes were washed three times for 5 min with TBST and incubated with HRP-conjugated secondary antibody of goat *anti*-rabbit IgG (1/10,000 dilution). The membranes were incubated for 1 h and then washed thrice for 5 min with TBST. The immunoreactive bands were revealed by chemiluminescence (ECL Western Blotting Substrate, Solarbio, Beijing, China) and measured by using ChemiScope 6000 (CliNX, Shanghai, China).

### Immunohistochemistry assay

The IHC assay was performed as previously described [26]. In brief, both the gill and muscle tissues were fixed with 4 % paraformaldehyde for 24 h at 4 °C, and then paraffin-embedded samples were cut into 4-μm sections and baked at 60 °C for 2 h. Sections adhered to slides were deparaffinized with xylene and rehydrated, submerged into EDTA antigenic retrieval buffer and microwaved for antigenic retrieval for 15 min. Later, the sections were treated with 3 % hydrogen peroxide (H_2_O_2_) in methanol, followed by incubation with 1 % bovine serum albumin (BSA) to block nonspecific binding at RT for 1 h. Then the tissue sections were incubated with the primary antibody rabbit *anti*-*Mr*-GS (1/200 dilution) overnight at 4 °C. After washing thrice with TBST, slides were incubated with HRP-conjugated goat *anti*-rabbit IgG (1/1000 dilution) at RT for 1 h and developed with DAB substrate solution (Guge Biotech, China). Finally, the sections were counterstained with hematoxylin, mounted and photographed (ECLIPSE E100, Nikon).

### Determination of Glutamine concentration

Frozen gills and muscle tissues were processed and measured using shrimp Glutamine ELISA Kit (Kawanshu, Shanghai, China) following manufacturer instruction. Briefly, to the microwells previously coated with *anti*-Gln antibodies, samples, standards, and HRP-labeled detection antibodies were sequentially added with appropriate incubations and washing. Tetramethylbenzidine (TMB) was added to each microplate wells to form a final yellow color from blue by the catalysis of peroxidase. The absorbance was measured at a wavelength of 450 nm using a microplate reader (Molecular Devices, USA), the intensity of color was measured.

## Results

### Sequence and phylogenetic analysis of *Mr*-GS

The full-length cDNA transcript of *Mr*-GS was 1965 bp with a 76 bp at 5′-untranslated region (UTR), an 803 bp 3′-UTR containing a 13 bp poly (A) tail. Nucleotide sequence analysis showed that the ORF of 1086 bp which encoded a putative protein of 361 AA with estimated molecular weight (MW) of 40.75 kDa (Fig 1A). SMART analysis displayed that *Mr*-GS protein contained two catalytic domains of Gln-synt_N located at N-terminal region at about 21-101 bp and Gln-synt_C located at C-terminal region at 107-356 bp (Fig 1B).

**Fig 1.**
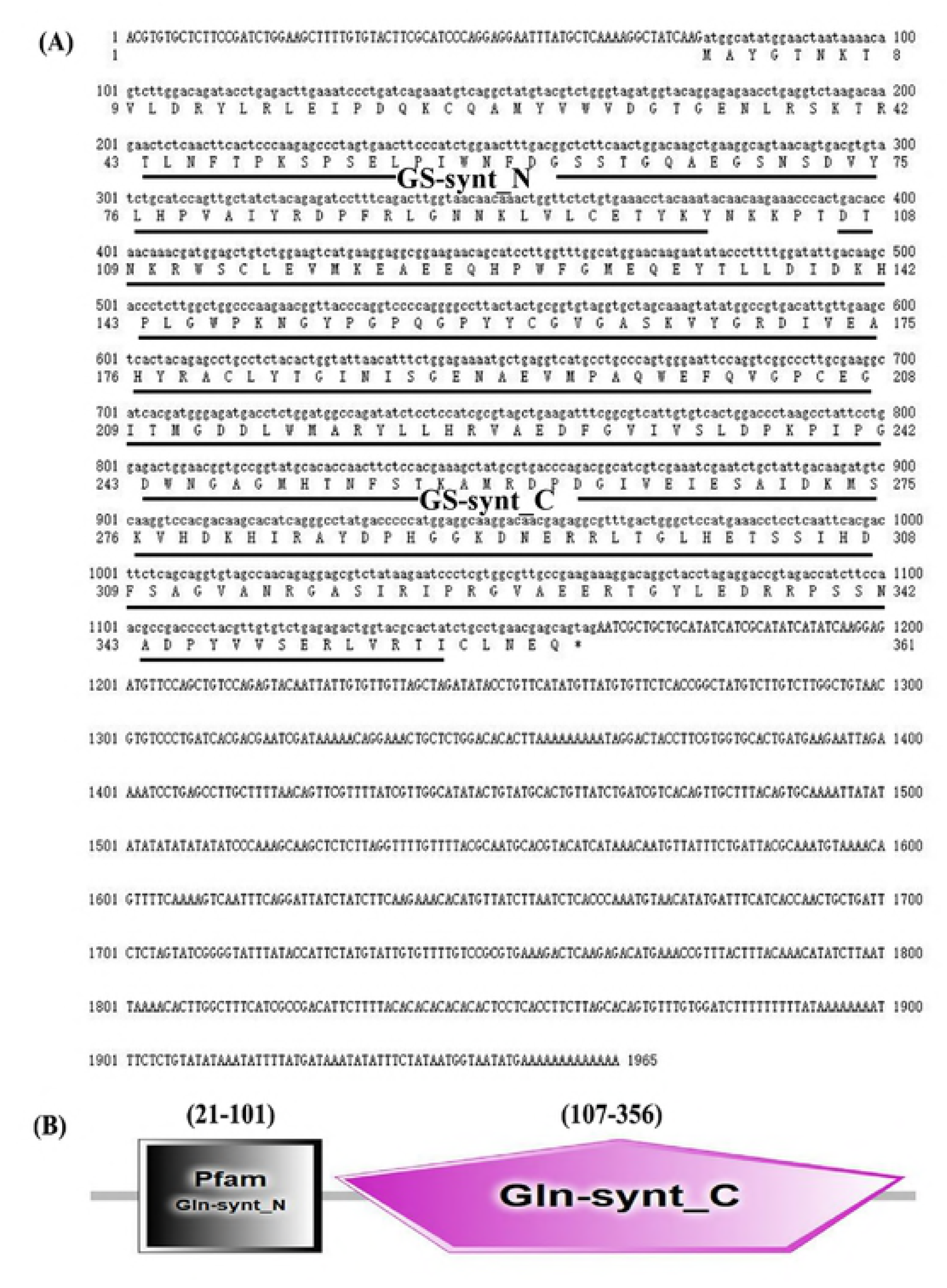
The sequence analysis of *Mr*-GS. (A) The full-length cDNA sequence and deduced amino acid sequence of *Mr*-GS. The ORF of the nucleotide sequence was shown in lowercase, while the 5’ and 3’-UTR sequences were shown in upper-case letters. The two potential *Mr*-GS binding domains were underlined in black. (B) Architecture and location representation of two characteristic domains of *Mr*-GS.

Multiple sequence alignment indicated that the AA ORF sequences of *Mr*-GS showed 87 % identity to the *Marsupenaeus japonicus* GS (*Mj*-GS), 83 % to *Hyalella azteca* GS (*Ha*-GS), 81 % to *Pacifastacus leniusculus* GS (*Pl*-GS), 79 % to *Daphnia magna* GS (*Dm*-GS), 71 % to *Danio rerio* GS (*Dr*-GS), and 70 % identity to the *Homo sapiens* GS (*Hs*-GS) (Fig 2). The results suggested that GS proteins were highly conserved from invertebrates to vertebrates, and multiple alignments discovered five conserved regions within *Mr*-GS. Besides, the phylogenetic tree demonstrated that GS genes were separated into two groups consisting of invertebrate and vertebrate. The invertebrates included crustacean, insecta, arachnida, and merostomata, and the vertebrates contained actinopterygii and mammalians, correlating well with the evolutionary origins. As shown in Fig 3, the GS of *M*. *rosenbergii* were clustered with *Litopenaeus vannamei*, *Marsupenaeus japonicus*, *Fenneropenaeus chinensis*, *Penaeus monodon*, *Hyalella azteca*, *Procambarus clarkii*, *Pacifastacus leniusculus* and *Daphnia magna* together into the crustacean group.

**Fig 2.** Analysis of protein sequences across species. Five conserved regions were marked with I: the latch (F/Y-D-G-S-S), II: (G-X(8)-E/K-V-X(3)-Q-W-E), III: ATP-binding site (K-P-X(4,5)-N-G-A-G-X-H-T-H-T-N-X-S), IV: Glutamate binding site (N/S-R-X(3)-I-R-I-P-R), and V: (F/L-E-D-R-X-P-S-X-N-X-D-P-Y), respectively. Multiple-sequence alignment of *M*. *rosenbergii* (*Mr*-GS) with Dr-GS, *Danio rerio* GS (NP 878286.3); Dm-GS, *Daphnia magna* GS (KZS15608.1); Ha-GS, *Hyalella azteca* GS (XP 018023652.1); Pl-GS, *Pacifastacus leniusculus* GS (AFV39702.1); Hs-GS, *Homo sapiens* GS (NP 001028216.1); Mj-GS, *Marsupenaeus japonicus* GS (AWW43688.1).

**Fig 3.** Phylogenetic analysis of full-length amino acid sequences of *Mr*-GS. The diagram was generated by the neighbor-joining method using the MEGA 6.0 program. Numbers next to the branches represent the percentage of replicate trees in the bootstrap replication (1,000). Bar scale at the bottom indicates 5 % amino acid divergence. The analysis involved 20 amino acid sequences including *Macrobrachium rosenbergii* (marked with a triangle); *Litopenaeus vannamei* (AEO80035.1); *Marsupenaeus japonicus* (AWW43688.1); *Fenneropenaeus chinensis* (AFN66649.1); *Penaeus monodon* (AGA83299.1); *Hyalella azteca* (XP 018023652.1); *Procambarus clarkii* (AKN79748.1); *Pacifastacus leniusculus* (AFV39702.1); *Daphnia magna* (KZS15608.1); *Halyomorpha halys* (XP 014294596.1); *Leptinotarsa decemlineata* (XP 023022227.1); *Anoplophora glabripennis* (XP 018567831.1); *Anopheles darlingi* (ETN61772.1); *Centruroides sculpturatus* (XP 023214668.1); *Limulus polyphemus* (XP 013778191.1); *Parasteatoda tepidariorum* (XP 015929268.1); *Danio rerio* (NP 878286.3); *Homo sapiens* (NP 001028216.1); *Bos taurus* (AAI03100.1); *Mus musculus* (AAA37746.1); *Rattus norvegicus* (NP 058769.4).

### Tissue distribution and mRNA expression profiles of Mr-GS

GS gene expression in the control group (non-stress treated prawn) was analyzed by qRT-PCR. The level of the *Mr*-GS mRNA could be detected in all tested tissues, organized from high to low expression levels as muscle> gills> heart> stomach> brain > haemolymph (Fig 4).

**Fig 4.** Tissue distribution analysis of *Mr*-GS in non-salinity stress of *M*. *rosenbergii* by qRT-PCR. Relative expression levels of *Mr*-GS in the gill, muscle, stomach, heart, brain, and hemolymph, β-actin as an internal control gene.

In response to hyperosmotic stress (0→13 ‰), the expression of the mRNA levels were found to be increasingly up-regulated both in the gills and muscles in a timely manner. Compared to that of 0 h, the mRNA expression was significantly increased in the gills with 2.54, 4.21, and 10.83 folds at 3, 6 and 12 h post of the treatment (Pt). While it was also considerably increased in the muscles with 11.66, 17.97, and 45.92 folds at 3, 6 and 12 h Pt (Fig 5A). On the other hand, under the hypo-osmotic stress (13→0 ‰), compared to that of 0 h, the mRNA expression was significantly increased in the gills with 1.63, 3.30, and 3.52 folds at 3, 6 and 12 h Pt. While it was also increased dramatically in the muscles with 4.06, 42.99, and 26.69 folds at 3, 6 and 12 h Pt (Fig 6 A).

**Fig 5.** Expression levels of *Mr*-GS exposed to hyperosmotic stress. (A) Relative mRNA expression levels of *Mr*-GS exposed to hyperosmotic stress at 0, 3, 6, and 12 h in the gill and muscle, with β-actin as an internal gene. These results are means ± SD. \**p*<0.05, \*\**p*<0.01 versus control. (B) Relative expression levels of *Mr*-GS protein levels exposed to hyperosmotic stress in gill and muscle, detected by western blot analysis, β-actin as control, and immunohistochemistry of gill and muscle tissues at 0 and 12 h. Bars: 5 μm.

**Fig 6.** Expression levels of *Mr*-GS exposed to hypoosmotic stress. (A) Relative mRNA expression levels of *Mr*-GS exposed to hypoosmotic stress at 0 h, 3 h, 6 h and 12 h in the gill and muscle, with β-actin as an internal gene. These results are means ± SD. \**p*<0.05, \*\**p*<0.01 versus control, ns = non-significant. (B) Relative expression levels of *Mr*-GS protein levels exposed to hyperosmotic stress in gill and muscle, detected by western blot analysis, β-actin as control, and immunohistochemistry of gill and muscle tissues at 0 and 12 h. Bars: 5 μm.

### Western blotting and Immunohistochemistry analyses

To further investigate the function of GS protein in osmoregulation in *M*. *rosenbergii*, we evaluated the GS protein expression levels in gill and muscle tissues. Compared with that at 0 h, the results showed that the expression of *Mr*-GS protein was considerably up-regulated in both the gills and muscle at 12 h Pt (Fig 5B, 6B). Furthermore, the same tissue samples were subjected to the IHC assay. The results showed a similar tendency of protein expression levels with that of the western blotting in both the muscle and gill tissues (Fig 5B, 6B).

### Changes of Gln concentration

There have been shown that Gln played a crucial role in osmoregulation; therefore, we analyzed the Gln concentration in the gill and muscle of the prawns. Compared to that of 0 h, in response to hyperosmotic stress, Gln concentration was almost increased in both gills and muscle at 3, 6 and 12 h post treatment with 0.49, 1.83, 2.02 folds, and 1.16, 1.41, 1.29 folds, respectively. While, under hypo-osmotic stress, Gln concentration was increased in both gills and muscle at 3, 6 and 12 h post treatment with 3.99, 3.40, 2.59 folds, and 1.72, 1.83, 1.80 folds respectively (Fig 7).

**Fig 7.** Change folds of Glutamine concentrations in the *M*. *rosenbergii* under hyperosmotic stress. (A) and hypoosmotic stress (B) at 3, 6 and 12 h in the gills and muscles. These results are means ± SD.

## Discussion

In the present study, we firstly cloned the glutamine synthetase (*Mr*-GS) of *M*. *rosenbergii*, then evaluated its roles in osmoregulation under osmotic stress. There are two catalytic domains, named Gln-synt_N and Gln-synt_C in *Mr*-GS, which are essential for the activity of GS. Multiple sequence alignment indicated that *Mr*-GS proteins were highly conserved with five conserved regions that are present from invertebrates to vertebrates [18]. Up to date, there are three types of the cytosolic GS gene. GS I and III located mostly in prokaryotes, and GS II was identified from eukaryotes [12, 27]. In this report, *Mr*-GS belonged to the GS II. A phylogenetic tree showed that it has a close association with other crustacean groups GS gene. Besides, in the present study, the GS proteins were separated into six clades including the crustaceans, insecta, arachnida, merostomata, actinopterygii, and mammalian groups, which was consistently correlated with the evolutionary origin of GS.

The mRNA transcripts of *Mr*-GS could be detected in all examined tissues and indicated that GS is a widely distributed enzyme, this was also observed in other species such as *Litopenaeus vannamei* [4] and *Fenneropenaeus chinensis* [12]. There has been reported that more energy was required to maintain the body metabolic balance and osmoregulation when the shrimps were exposed to salinity stress [28, 29]. It is also stated that crustaceans increase their oxygen consumption, respiratory quotient and enhance the protein catabolism rate to maintain energy consumption to resist environmental stresses [18, 28, 29]. Muscle tissue is the largest storehouse of protein and AA in prawns [14]. During the stress, due to the more active catabolism of protein and AA, the ammonia concentration usually raised in the tissues, resulting in the toxicity to the host. Therefore, the extra ammonia must be secreted to the water or converted to glutamine [18, 21]. Our results indicated that during the stress exposure, *Mr*-GS expression levels had significantly increased in the gills and muscles at various time points. This might relate to the conversion of ammonia which is catalyzed by GS. Previous reports proposed that the ammonia formed during catalysis of AA in aquatic crustacean is mainly transported to gills and excreted as free ammonia by diffusion [30, 31]. Increased expression of GS in the gills and digestive tissues of *C*. *gigas*, and the protein levels of GS were significantly regulated in the muscles of *Monopterus albus* when exposed to stress conditions [17, 25]. In this report, the ammonia concentration in the prawn was not measured, which needs to be investigated in the future. Similar studies have reported that FAA function as an essential osmoregulators in crustaceans such as *Panopeus herbstii* [32], *P*. *monodon* [33] and *M*. *nipponense* [20]. In *M*. *rosenbergii*, the total FAA including glycine, proline, arginine, glutamate, and alanine concentrations were maintained nearly 1 mM in freshwater; however, it dramatically increased up to 2.1 mM in higher salinities [19]. Gln has been regarded as one of the essential osmolytes in *P*. *motoro* [34], *L*. *vannamei* [18] and *M*. *albus* [25]. In summary, Gln not only performing as a nontoxic transporter of ammonia but also a kind of abundant FAA and as a major osmolyte for crustaceans.

## Conclusion

In conclusion, our results revealed significant evidence that *Mr*-GS could be involved in coordinate osmoregulation in *M*. *rosenbergii* exposed to osmotic stress. Our results will shed new light on the osmoregulation of crustacean.

## Acknowledgments

This work was jointly supported by “Innovation and Strong Universities” special fund from the Department of Education of Guangdong Province (KA170500G, TK222001G, KA18058B3, KA1819604). Fund from the Department of Science and Technology of Guangdong Province (KA1810312), V Sarath Babu was supported by Chinese Postdoctoral Science Foundation (189103).

## Author Contributions

**Conceived and designed the experiments:** Li Lin, Gan Pan. **Performed the experiments:** Zhijie Lu. **Analyzed the data:** Zhendong Qin, Chengkai Ye, Jiabo Li, Sarath Babu V, Guang Yang, Haiyang Shen, Guomao Su. **Writing:** Zhijie Lu, Sarath Babu V, Li Lin.

